# A global map of RNA binding protein occupancy guides functional dissection of post-transcriptional regulation of the T cell transcriptome

**DOI:** 10.1101/448654

**Authors:** Adam J. Litterman, Wandi S. Zhu, Robin Kageyama, Wenxue Zhao, Noah Zaitlen, David J. Erle, K. Mark Ansel

## Abstract

RNA binding proteins (RBPs) mediate constitutive RNA metabolism and gene specific regulatory interactions. To identify RNA cis-regulatory elements, we developed GCLiPP, a biochemical technique for detecting RBP occupancy transcriptome-wide. GCLiPP sequence tags corresponded with known RBP binding sites, specifically correlating to abundant cytosolic RBPs. To demonstrate the utility of our occupancy profiles, we performed functional dissection of 3′ UTRs with CRISPR/Cas9 genome editing. Two RBP occupied sites in the CD69 3′ UTR destabilized the transcript of this key regulator of lymphocyte tissue egress. Comparing human Jurkat T cells and mouse primary T cells uncovered hundreds of biochemically shared peaks of GCLiPP signal across homologous regions of human and mouse 3′ UTRs, including a cis-regulatory element that governs the stability of the mRNA that encodes the proto-oncogene PIM3 in both species. Our GCLiPP datasets provide a rich resource for investigation of post-transcriptional regulation in the immune system.

## Introduction

The life cycle of protein coding RNA transcripts involves their transcription from DNA, 5′ capping, splicing, 3′ polyadenylation, nuclear export, targeting to the correct cellular compartment, translation and degradation (Beelman and Parker, 1995; Martin and Ephrussi, 2009; Reed, 2003). RNA binding proteins (RBPs) coordinately regulate these processes through interaction with RNA cis-regulatory elements, often in the 5′ and 3′ untranslated regions (UTRs) whose sequences are not constrained by a functional coding sequence (Keene, 2007). Mammalian genomes encode hundreds of RBPs (Castello et al., 2012), and mutations in individual RBPs or even individual binding sites can induce strong developmental, autoimmune and neurological defects in human patients and mouse models (Bassell and Kelic, 2004; Kafasla et al., 2014; Schwerk and Savan, 2015). As much as half of the extensive gene expression changes that occur during T cell activation occur post-transcriptionally (Raghavan et al., 2002), and several RBPs are known to be critical determinants of immune function and homeostasis (Kafasla et al., 2014).

Methods like DNase I hypersensitivity and ATAC-seq that query regulatory element accessibility and occupancy without prior knowledge of their protein binding partners have proven themselves as powerful techniques for the systematic mapping of *cis*-regulatory sequences in DNA (Buenrostro et al., 2013; Thurman et al., 2012). Their development has allowed for comparisons in the regulatory structure of diverse cell types (Corces et al., 2016) and across the tree of life (Villar et al., 2015; Wilson et al., 2008). A lack of analogous systematic methods for mapping the transcriptome’s cis-regulatory landscape has limited our understanding of post-transcriptional regulatory circuits and the evolution of untranslated regions of transcribed genes.

Current methods for regulatory element identification in RNA have focused on specific *trans* factors (Lee and Ule, 2018), although more recent technologies have also analyzed secondary structure (Spitale et al., 2015) and interaction with chromatin (Li et al., 2017) transcriptome wide. Protein precipitation (Baltz et al., 2012) and chemical biotinylation of proteins (Freeberg et al., 2013) have been used to analyze global RBP occupancy in cell lines and yeast, respectively, but difficulty remains in defining RNA regulatory activity in a systematic way. Here, we create global RBP occupancy maps for a human T cell line, Jurkat, and primary mouse T cells. Comparing RBP occupancy for thousands of mRNAs across species identified biochemically shared regulatory sites, which are enriched for phylogenetically conserved sequences. Finally, we used a scalable system of CRISPR dissection to define regions of functional activity in 3′ UTRs of mouse and human transcripts of immunological importance. Biochemically derived maps of RBP occupancy are a powerful tool for the interrogation of post-transcriptional gene regulation in the immune system.

## Results

### Transcriptome-wide analysis of RBP occupancy in T cells in two species

To achieve transcriptome-wide RBP binding site profiling, we developed a protocol for Global Cross-linking Protein Purification (GCLiPP) suitable for use in mammalian cells and applied this technique in human Jurkat T cells and cultured primary mouse T cells (Figure 1A). GCLiPP is an adaptation of previously described biochemical methods for crosslinking purification of all mRNA-RBP complexes. The key features of GCLiPP include: crosslinking of endogenous ribonucleoprotein complexes using high energy UV light (no photo-crosslinkable ribonucleotide analogues); oligo-dT pulldown prior to biotinylation to enrich for mRNA species; chemical biotinylation of primary amines using a water soluble reagent with a long, flexible linker; brief RNase digestion with RNase T1; and on-bead linker ligation with radiolabeled 3′ linker to facilitate downstream detection of ligated products. We used the guanine specific ribonuclease T1 to favor larger average fragment sizes than using an RNA endonuclease with no nucleotide specificity (such as RNase A) and ligated RBP protected fragments into a small RNA sequencing library.

**Figure 1.**
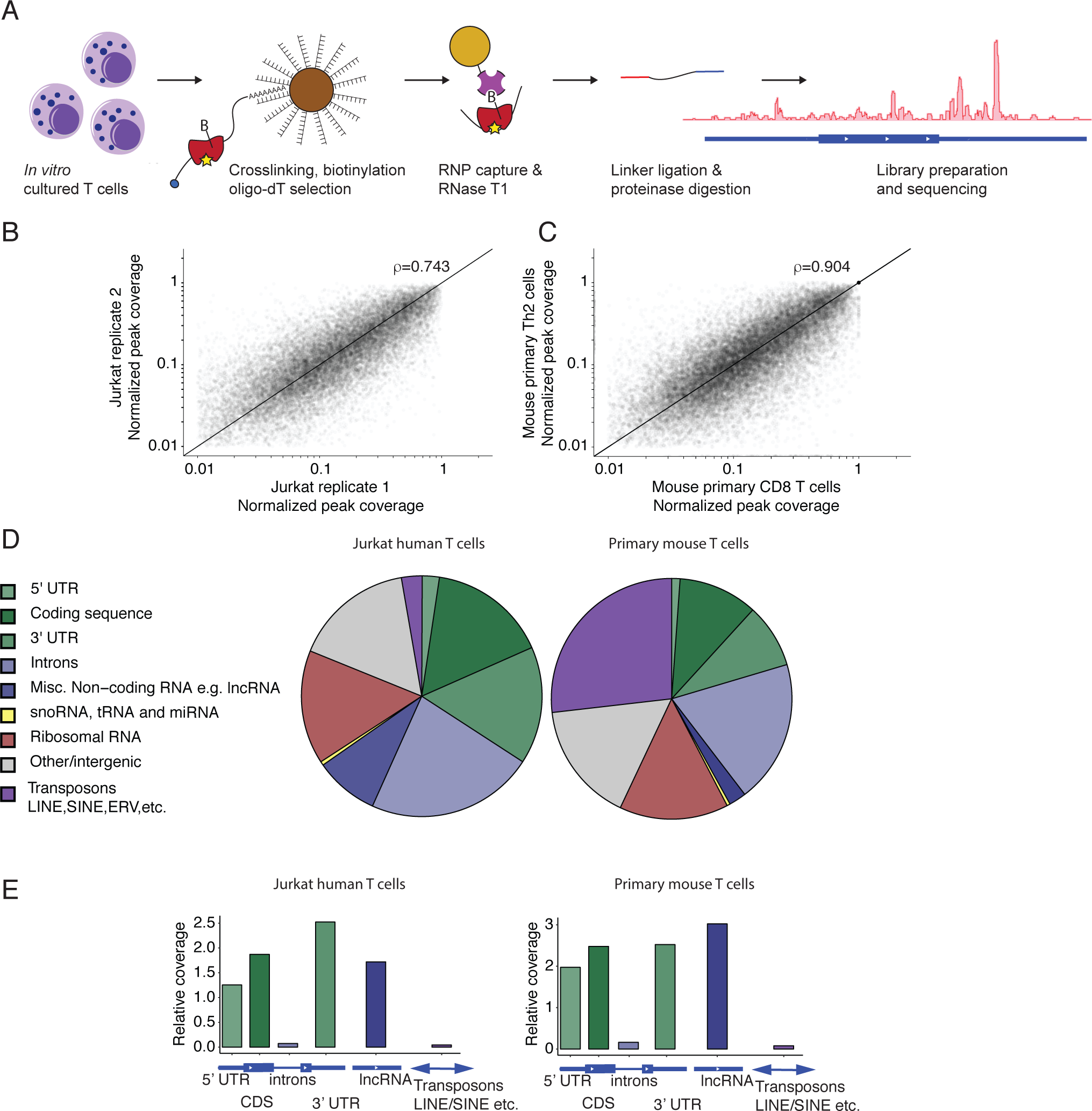
GCLiPP sequencing reveals RNA transcript protein occupancy. (**A**) GCLiPP method of global RBP profiling. T cell RNAs are crosslinked to RBPs and lysates are biotinylated on primary amines. mRNAs are enriched with oligo-dT beads, and RBP protected sites are digested, captured, sequenced and aligned to the genome. (**B**) Normalized GCLiPP read depth (fraction of reads in called peak relative to all GCLiPP reads in annotated 3′ UTR) in two replicates of Jurkat cells. r represents Pearson correlation. (**C**) Normalized GCLiPP read depth in mouse primary Th2 and CD8 T cells. r represents Pearson correlation. (**D**) Proportion of mapped GCLiPP reads derived from genomic features. (**E**) Relative coverage of genomic features in GCliPP sequencing reads relative to total length of genomic features of indicated class.

We called local peaks of GCLiPP sequence read density and measured the distribution of GCLiPP reads within those peaks to assess the reproducibility of the technique. Local read density within individual transcripts was similar between experiments, as GCLiPP fragments yielded highly reproducible patterns in technical replicates (e.g. comparing replicate Jurkat T cell samples, Figure 1B) and across multiple pooled experiments (e.g. comparing CD4^+^ and CD8^+^ T cells, Figure 1C). A similar distribution of transcriptome features constituted GCLiPP libraries from both Jurkat and primary T cells, with read coverage strongly enriched within mature mRNAs and long non-coding RNAs (Figure 1D, E). The most striking difference was the greater proportion of reads derived from transposable elements in mouse GCLiPP libraries. This increase is likely due to the greater amount of annotated transposable elements in the mouse genome since the relative coverage of these elements was similar between species.

### GCLiPP read density represents cytosolic RBP occupancy

To validate GCLiPP, we systematically examined the relationship between GCLiPP occupancy profiles in human Jurkat cells and enhanced cross-linking immunoprecipitation (eCLIP) analyses of specific RBP binding profiles in K562 cells from the Encyclopedia of DNA Regulatory Elements (ENCODE) project (Sundararaman et al., 2016). We examined pairwise correlations of normalized read density across individual 3′ UTRs between GCLiPP and individual RBP eCLIP samples to identify contributions of each RBP, and also compared GCLiPP and the input control for each eCLIP experiment (Figure 2A). eCLIP for many RBPs, such as TIA1 and IGF2BP1, more closely matched GCLiPP read density than the eCLIP control input across the genome (Figure 2B). We also found RBPs, such as PUM2, that exhibited anti-correlations with GCLiPP signal across individual transcripts. In these cases, we typically saw focal RBP binding to specific sites within transcripts (such as UGUA motifs in the case of PUM2) that, while represented in GCLiPP reads, did not dominate the GCLiPP signal (Figure 2A, bottom panel). Although the PUM2 eCLIP profile did not correlate to GCLiPP signal genome wide, PUM2 binding sites were still overrepresented in GCLiPP data. This was revealed when we called GCLiPP peaks with CLIPper (Lovci et al., 2013) and compared these peaks with CLIPper called peaks in eCLIP datasets. The observed fraction of PUM2 eCLIP peaks that overlap GCLiPP peaks (0.56) was much greater than the fraction overlapping eCLIP peaks randomly shuffled across the 3′ UTRs from which they were derived (Figure 2C, bottom panel). Similar results were obtained for TIA-1 (Figure 2C, top panel) and IGF2BP1 (Figure 2C, middle panel). These enrichments above background binding for IGF2BP1, TIA1 and PUM2 were amongst the highest 8 of the 87 RBPs whose eCLIP signals were examined (Supplementary Figure 1).

**Figure 2.**
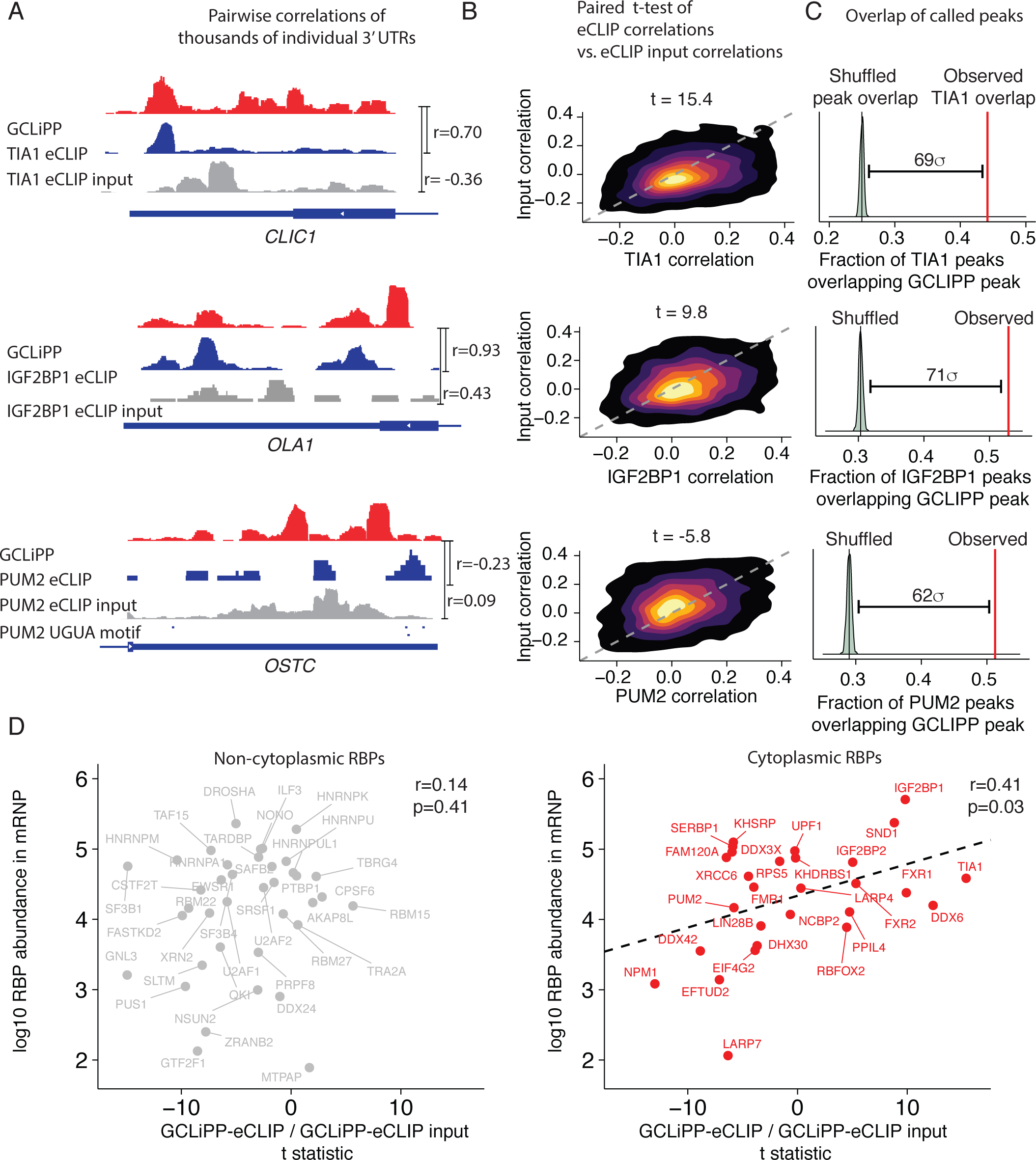
Comparisons with eCLIP reveal abundant cytosolic RBPs drive GCLiPP signal. (**A**) Genomic snapshots of individual 3′ UTRs showing exemplary correlation between eCLIP datasets and GCLiPP. GCLiPP is shown in red, while the indicated RBP eCLIP data is shown in blue, and matched control input samples are shown in gray, shown for the 3′ UTRs of the indicated gene. r indicates Pearson correlation between pairs of normalized read density at a given nucleotide for the indicated comparisons. (**B**) 2D density plots showing matched correlations between GCLiPP and eCLIP for the indicated RBP (X-axis) and GCLiPP and the matched control input sample (Y-axis) for individual 3′ UTR for all expressed genes in eCLIP and GCLiPP datasets. The t-statistic shown is for a paired t-test of the correlations. (**C**) Overlap of CLIPper called peaks in 3′ UTRs in GCLiPP and eCLIP. Red lines indicate observed overlap of GCLiPP peaks and eCLIP peaks. Green distribution represents bootstrapped expected overlap, computed by shuffling called eCLIP peaks within the same 3′ UTR, computing overlap of shuffled set with GCLiPP called peaks, and repeating this analysis 500 times. The indicated distance represents the number of standard deviations above the mean shuffled overlap of the observed overlap. (**D**) Correlation of eCLIP-GCLiPP paired t-tests (from (**B**) and RBP abundance in mRNPs). RBPs shown in gray score are not cytosolic localized (<5 cytosolic according to COMPARTMENTS) whereas RBPs in red are cytosolic localized (5 cytosolic according to COMPARTMENTS).

We performed genome wide correlation analysis for 87 RBPs obtained from eCLIP data, and compared the correlation between eCLIP and GCLiPP with RBP abundance previously determined via mass spectrometry (Baltz et al., 2012). There was an overall significant correlation between RBP abundance and correspondence between RBP eCLIP and GCLiPP profiles (r=0.28, p=.022). However, stratifying RBPs by their predominant cellular localization (Binder et al., 2014) showed that this correlation was driven almost entirely by cytosolic RBPs (Figure 2D). The fraction of eCLIP peaks that overlapped GCLiPP peaks above a shuffled background was also significantly greater for cytosolic versus non-cytosolic RBPs (p=0.003, Supplementary Figure 1 inset). These findings were expected, as the GCLiPP experimental protocol preferentially samples the cytosol by eliminating most nuclear material. In summary, we conclude that GCLiPP read density reflects transcriptome-wide cytosolic RBP occupancy.

### RBP Occupancy of known RNA cis-regulatory elements in primary T cells

We examined the GCLiPP profiles at previously characterized cis-regulatory elements of various functional and structural categories in primary mouse T cells. The canonical polyadenylation signal AAUAAA is a known linear sequence motif that binds to a number of RBPs in the polyadenylation complex, including CPSF and PABP (Millevoi and Vagner, 2009), as part of constitutive mRNA metabolism. We examined T cell lineage-defining transcripts with well-resolved GCLiPP profiles (due to their high expression levels), including *Cd3g* (Figure 3A), *Cd3e, Cd4*, and *Cd8b1* (Supplementary Figure 2). The only canonical polyadenylation signal sequences in these transcripts were contained within called GCLiPP peaks, often as the peak with the highest GCLiPP read density in the entire transcript. Interestingly, the GCLiPP profile of *Cd8b1* contained direct biochemical evidence for alternative polyadenylation signal usage (Figure S2C), a phenomenon that has previously been described to be important in activated T cells (Sandberg et al., 2008). GCLiPP peaks appeared in multiple canonical polyadenylation signal sequences in *Cd8b1*, coincident with clear evidence for both short and long 3′ UTR isoform usage indicated by lower RNAseq read counts after the initial canonical polyadenylation signal. A similar pattern was also apparent in *Hif1a* (Figure S1D) and a number of other highly expressed transcripts.

**Figure 3.**
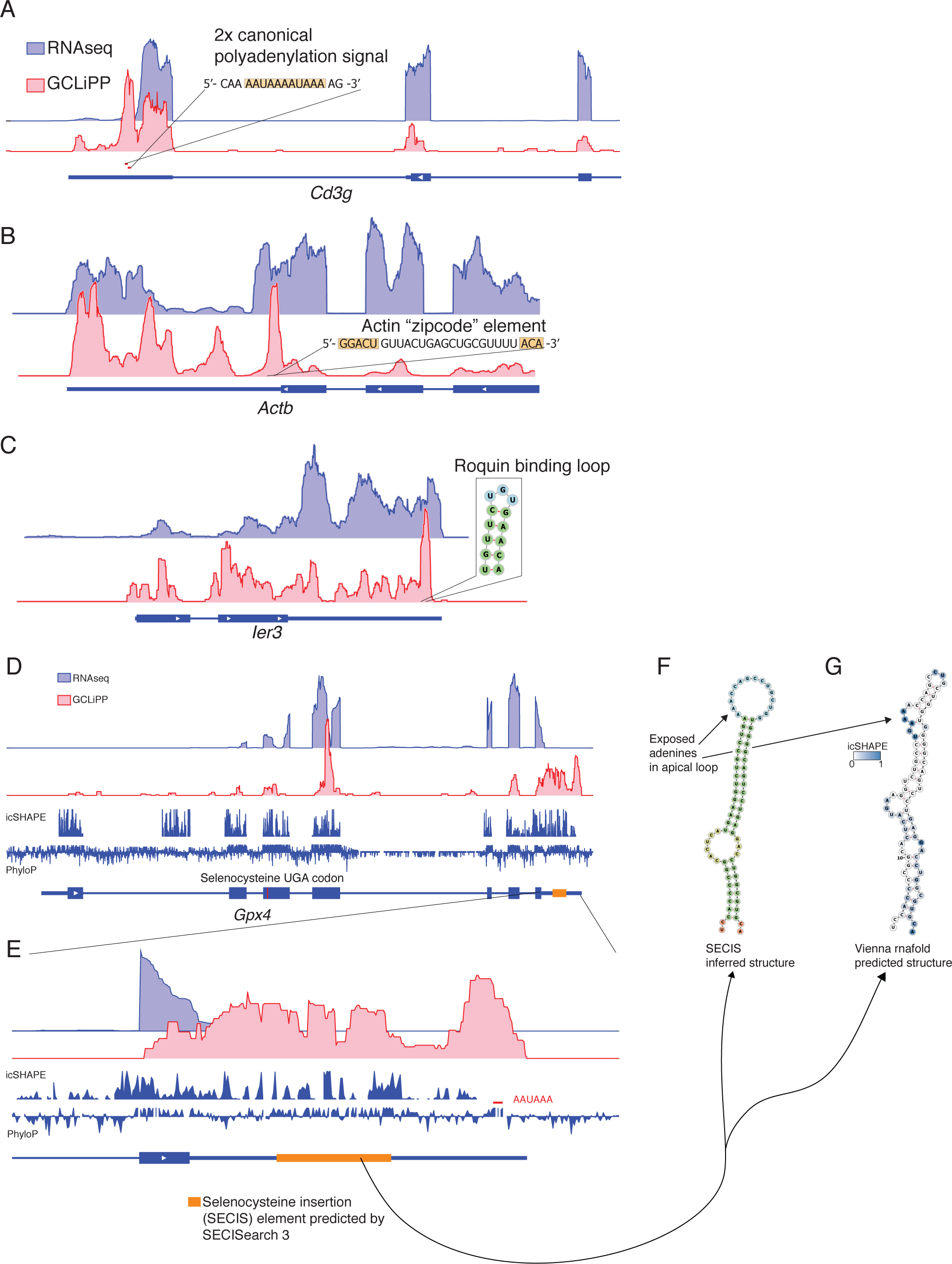
GCLiPP recapitulates previously described mRNA-RBP interactions in primary T cells. RNAseq and GCLiPP tracks for **(A)** *Cd3g* (**B**) *Actb* **(C)** *Ier3* and (**D-G**) *Gpx4*. RNAseq track is from resting Th2 cells. GCLiPP is sum of five experiments, three in Th2 and two in CD8 T cells. Location of known RBP binding determinants are shown as insets.

Known cis-regulatory elements involved in transcript localization were also represented by local regions of GCLiPP read density. The Beta-actin “zipcode” element is responsible for localization of *Actb* mRNA to the cellular leading edge in chicken embryo fibroblasts (Kislauskis et al., 1994) and contains conserved linear sequence elements separated by a variable linker. These conserved sequence elements are thought to form the RNA/protein contacts in a complex involving the actin mRNA and the RNA binding protein Igf2bp1 (previously known as Zbp1) where the non-conserved sequence winds around the RBP (Chao et al., 2010). This sequence corresponds to the center of the second highest peak of GCLiPP read density in the *Actb* transcript (Figure 3B). Some RBPs regulate the half-life and/or translation of the mRNAs that they bind. The mRNA-destabilizing Roquin/Regnase binding site in the 3′ UTR of *Ier3* is a straightforward example of this functional category of RNA/RBP interaction detected as a region of GCLiPP read density (Figure 3C).

The insertion of the selenium containing amino acid selenocysteine into selenoproteins represents a unique case of RBP regulation of protein translation. Selenoproteins are redox enzymes that use selenocysteine at key reactive residues (Johansson et al., 2005; Papp et al., 2007) Selenocysteine is encoded by the stop codon UGA, and this recoding occurs only in mRNAs that contain 3′ UTR *cis*-regulatory elements (termed SECIS elements) that bind to RBPs that recruit the elongation factor Eefsec and selenocysteine-tRNA (Berry et al., 1993; Tujebajeva et al., 2000). SECIS elements were prominent peaks of GCLiPP read coverage in selenoprotein mRNAs. For example, the predicted SECIS element (Mariotti et al., 2013) in the 3′ UTR of *Gpx4* was entirely covered by GCLiPP reads (Figure 3D). Indeed, a canonical polyadenylation signal and the full hairpin structure containing the SECIS element account for essentially all of the GCLiPP reads in the *Gpx4* 3′ UTR (Figure 3E). Comparing transcriptome-wide in vivo folding data from icSHAPE (Spitale et al., 2015) and GCLiPP data supports the identification of an RBP bound, structured SECIS element (Figure 3F,G). Furthermore, this analysis suggests that the folded, RBP bound structure is even larger than that predicted by SECISearch 3, with regions of GCLiPP read density and apposed high and low icSHAPE signals spanning almost the entire 3′ UTR. Thus, GCLiPP recapitulated previously described cis-regulatory elements that mediate constitutive RNA metabolism, transcript localization, regulation of gene expression, and translation, including both structured elements and single-stranded RNA determinants.

### GCLiPP-guided CRISPR dissection of immune gene post-transcriptional regulation

We then sought to use our GCLiPP RBP occupancy profiles to guide experimental dissection of the post-transcriptional regulation of immunologically important transcripts. We first focused on CD69, a cell surface C-type lectin protein transiently upregulated on T cells early during activation. CD69 inhibits lymphocyte egress from lymphoid organs, and has been implicated in a variety of other immune cell functions (Cibrián and Sánchez-Madrid, 2017). As *CD69* mRNA is a labile transcript (Santis et al., 1995) we sought to identify cis-regulatory elements in the 3′UTR that regulate stability. First, we designed guide RNAs (gRNA) targeting nucleotide positions 57 and 784 downstream of the stop codon to generate large deletions in the 3′UTR by transfecting CRISPR-Cas9 ribonucleoprotein complex (crRNP) in Jurkat cells. From the pool of transfected cells, we generated a clone (3′UTR Δ57-784) that contained two mutant alleles with deletions that spanned positions 22 to 853 and 44 to 833 of *CD69* 3′ UTR (Figure 4A). As a control, we generated a wildtype (WT) clone from Jurkat cells transfected with a scrambled control gRNA crRNP. Homozygous deletion of most of the *CD69*3′ UTR led to higher basal expression of CD69 protein (Figure 4B, left) and higher expression after stimulation with PMA and Ionomycin (Figure 4B, right). Importantly, the *CD69* transcript in the 3′UTR Δ57-784 clone decayed at a much slower rate, with a half-life of greater than 3 hours compared to 0.36 hours in WT Jurkat cells after global transcriptional inhibition with actinomycin D (Figure 4C). This effect was specific to *CD69* as the half-life of dual-specific phosphatase 2 (*DUSP2*), another labile transcript, was similar in WT and mutant clones (Figure 4C). These data indicate that the *CD69* 3′UTR contains destabilizing cis-regulatory elements responsible for the short half-life of the mRNA.

**Figure 4.**
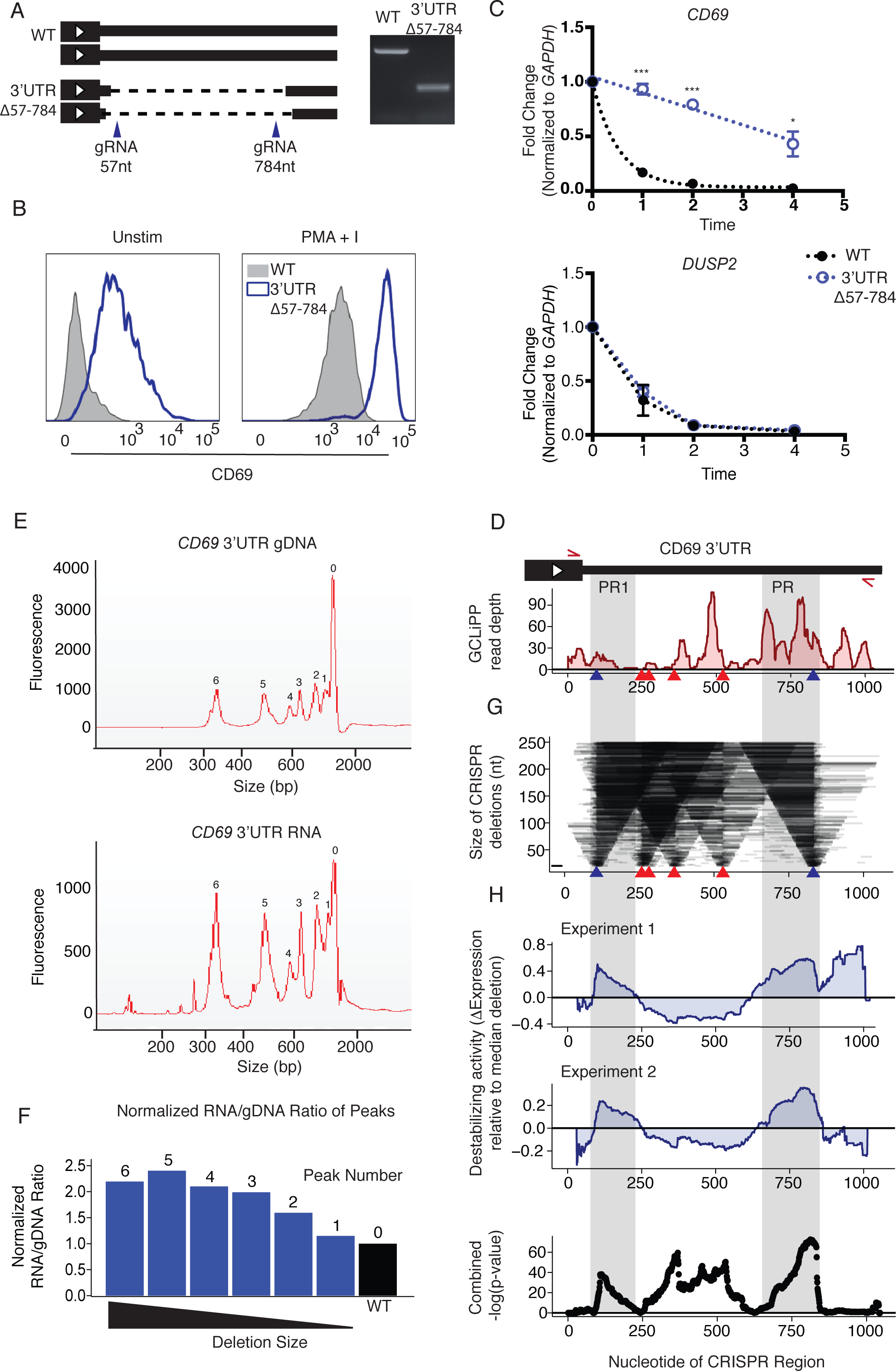
GCLiPP guides identification of destabilizing regions in CD69 3′UTR. (A) Schematic illustration and gel image of Jurkat WT and CD69 3′UTR edited clone (3′UTR Δ57-784). Editing was performed using CRISPR-Cas9 and verified through PCR. Blue arrows indicate gRNA placement with their positions in the 3′UTR indicated by nucleotide (nt) number. (B) CD69 protein expression of WT and 3′UTR Δ57-784 clone as measured by flow cytometry. Cells were either untreated or stimulated with PMA and Ionomycin (PMA + I) for 4 hours. (C) mRNA decay of *CD69* (top) and *DUSP2 (bottom)* transcript in WT and 3′UTR Δ57-784 clones. Cells were stimulated for 4 h with PMA+I and then treated with Actinomycin-D (act-D). Transcript expression was measured 0, 1, 2 or 4 hours post addition of act-D by qPCR and normalized to *GAPDH* expression. Data was generated from two separate experiments each with N=2 and significance was calculated using multiple t-test corrected with Holm-Sidak method *p<0.05, **p<0.01, ***p<0.001 (D-H) CRISPR-Cas9 dissection of *CD69* 3′UTR. (D) CD69 3′UTR GCLiPP peaks aligned to schematic illustration of 3′UTR. Red arrows indicate primer position during PCR amplification of gDNA and RNA from pooled crRNP transfected Jurkats. Arrow heads represent gRNA placement. Blue arrow heads were gRNAs also used in experiment in Figure 4A. (E) Microfluidic capillary electrophoresis of CD69 3′UTR gDNA (top) and reverse transcribed cDNA (bottom). Individual fragments identified by the Agilent Bioanalyzer software are indicated by the numbers on the graph. (F) Ratio of RNA/gDNA for each labeled peak in (D) normalized to predicted wild-type allele (black bar). Estimated molarity of each peak from the electrophoresis was used to calculate RNA/gDNA ratio. The labeled peaks are plotted in descending order based on deletion size. (G) Size of deletions generated using CRISPR-Cas9. Arrow heads represent gRNA placement as mentioned for Figure 4D. (H) Change in expression along the 3′UTR relative to median expression of all possible deletions. Per-nucleotide effect score was calculated by comparing median normalized RNA/gDNA ratio for all shown deletions spanning a given nucleotide with all shown deletions. Experiment 1 and 2 are duplicate samples which were transfected with 80µM or 120µM of gRNAs respectively. Grey shaded area PR1 and PR2 indicate regions of significant destabilizing activity. Unadjusted -log10 p-values from Welch’s two sample t-test comparing all deletions spanning a nucleotide with all other deletions across both experiments (bottom).

To determine whether RBP-occupied sites in the 3′UTR contain cis-regulatory elements that regulate stability, we performed CRISPR-Cas9 dissection of the region (Zhao et al., 2017). Using the GCLiPP profile as a guide, we designed 6 gRNAs along the 3′UTR, transfected them as a crRNP pool into Jurkat cells, and (RT)-PCR amplified the CD69 3′ UTR from genomic DNA and RNA from transfected cells (Figure 4D). The dissection led to many distinct short and long deletions (Figure 4E) that possessed destabilizing activity, indicated by a high RNA/gDNA ratio relative to the predicted WT allele as measured by microcapillary gel electrophoresis (Figure 4F). To determine whether certain RBP-occupied regions had greater destabilizing activity than others, we sequenced amplicon fragments to measure the relative abundance of transcripts containing the various deletions (Figure 4—source data 1), analyzed deletions <250bp and calculated relative RNA/gDNA ratios along the 3′UTR. Our analysis revealed varying deletion sizes (Figure 4G) and identified two regions with the highest destabilizing activity that correspond with GCLiPP peaks (protein occupied regions PR1 and PR2 in Figure 4D, G-H). This pattern was replicated in duplicate experiments using different crRNP concentrations (Figure 4H). Destabilizing activity was highly significantly concentrated in regions PR1 and PR2. These findings demonstrate that GCLiPP can provide useful hypothesis-generating data for identifying 3′UTR cis-regulatory elements that contribute to post-transcriptional gene regulation.

### Cross-species comparison of GCLiPP reveals patterns of biochemically shared post-transcriptional regulation

Next, we sought to compare RBP occupancy in mouse and human T cells. To do so, we performed Clustal Omega sequence alignments of thousands of human 3′ UTRs and their corresponding sequences in the mouse genome, and then designed an algorithm to identify correlated peaks of normalized GCLiPP read density along the aligned nucleotides (Figure 5A). Using this approach, we identified 1047 high-stringency biochemically shared GCLiPP peaks derived from 901 3′ UTRs (Supplementary table 1). As a class, biochemically shared peaks exhibited significantly higher sequence conservation than the full 3′ UTRs in which they reside (Figure 5B). The highly conserved, biochemically shared peak in *USP25* exemplifies this general pattern (Figure 5C, right panel). However, many biochemically shared peaks did not exhibit corresponding increases in local sequence conservation. For example, the *ARRB2* mRNA that encodes b-arrestin, another regulator of T cell migration in response to chemoattractant gradients (Fong et al., 2002), exhibited a common peak of RBP occupancy in Jurkat cells and primary mouse T cells that is roughly equally conserved as the rest of the 3′ UTR (Figure 5C, left panel).

**Figure 5.**
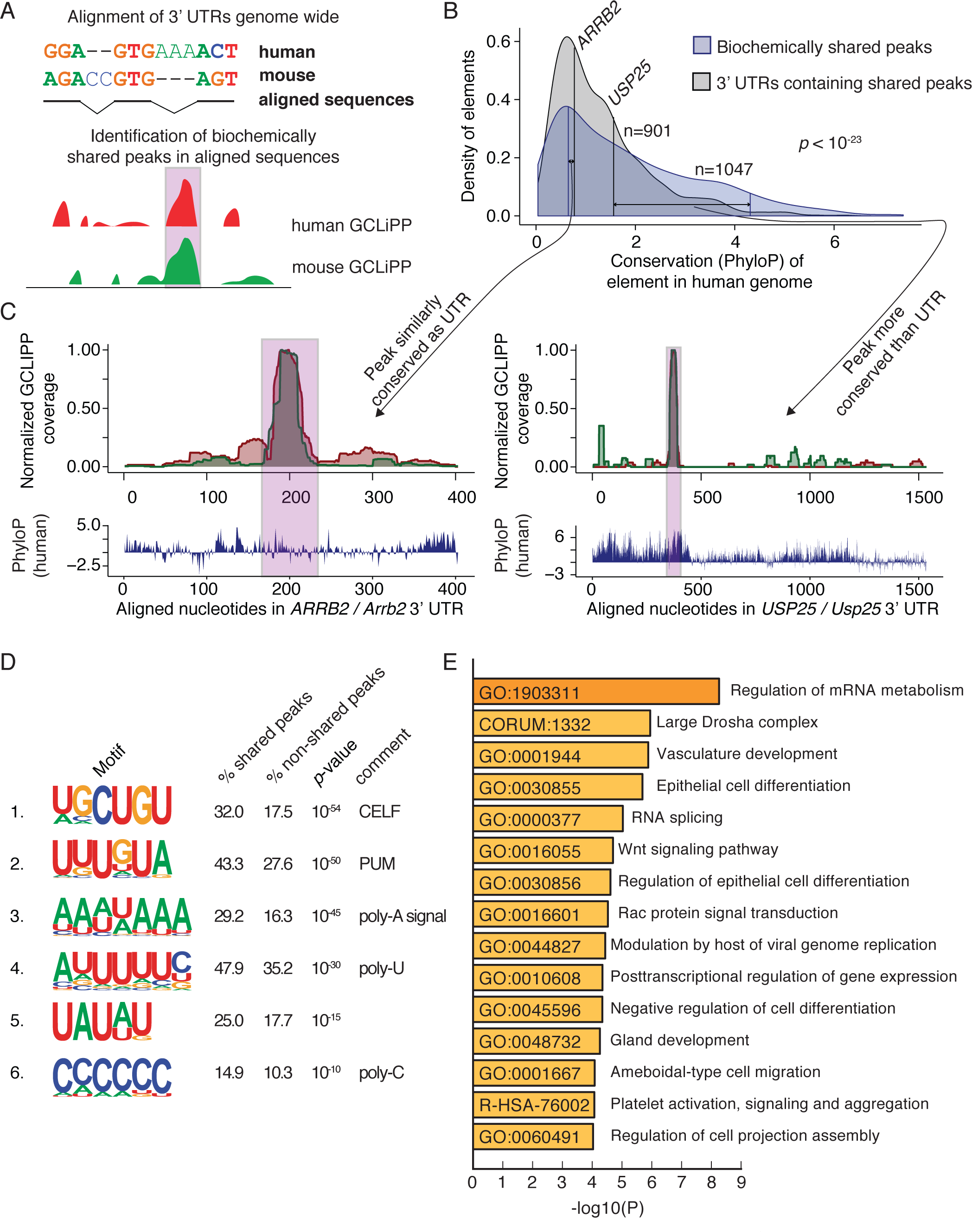
Comparison between mouse and human GCLiPP reveals principles of shared post-transcriptional regulation. (**A**) Schematic illustration of 3′ UTR alignment and biochemically shared GCLiPP peak calling. (**B**) Distribution of conservation across 100 vertebrates (PhyloP score) of regions in the human genome. Blue indicates biochemically shared peaks, gray indicates the 3′ UTRs of the transcripts that those peaks are contained within. For both peaks within *ARRB2* and *USP25*, their matched conservation of peak and UTR are indicated by connected vertical lines. (**C**) Human and mouse normalized GCLiPP density and conservation (PhyloP) across aligned nucleotides of the indicated 3′ UTRs. Biochemically shared peaks of GCLiPP read density are indicated in pink. (**D**) HOMER called motifs enriched in biochemically shared peaks. Percentages indicate the frequency of occurrence of the indicated motif in biochemically shared peaks versus non-shared background peaks. P-value indicates HOMER calculated p-value of enrichment. (**E**) Metascape called biological enrichment categories of genes containing biochemically shared peaks. The background set was all genes that contained peaks in both mouse and human GCLiPP datasets that did not contain a shared peak.

To examine which RBPs contributed to biochemically shared peaks more than other GCLiPP peaks, we used HOMER motif calling software (Heinz et al., 2010) to identify enriched motifs. Strikingly, of the six linear sequence motifs present in >10% of biochemically shared peaks with *p* ≤ 10^-10^, five resemble well-known regulatory sequences (Figure 5D). The two most common appeared to represent canonical CELF (Timchenko et al., 1996) and PUM (Hafner et al., 2010) binding motifs. Three other identified motifs corresponded to runs of homo-polymers: An A-rich motif that resembled the canonical polyadenylation signal (Proudfoot, 2011); a poly-U containing motif similar to a sequence that has long been known to stabilize mRNAs (Zubiaga et al., 1995) and a poly-C containing motif similar to the C-rich RNAs bound by poly-C binding proteins (Makeyev and Liebhaber, 2002). We used Metascape (Tripathi et al., 2015) to identify categories of biologically related genes enriched among mRNAs that contained biochemically shared GCLiPPpeaks (Figure 5E and Figure 5—source data 2). Interestingly, 3 of the 5 most enriched categories were related to RNA regulation (“regulation of mRNA metabolism,” “large Drosha complex,” “RNA splicing”), with the broad category “post-transcriptional regulation of gene expression” also in the top 10. Thus, biochemically shared GCLiPP binding sites are generally more well conserved than their local sequence context, are enriched for well-studied RBP binding motifs, and occur preferentially in genes that encode proteins involved in post-transcriptional gene regulation, suggestive of conserved autoregulatory gene expression networks.

We hypothesized that functionally conserved destabilizing cis-regulatory elements could be identified by examining biochemically shared GCLiPP peaks in 3′ UTRs of labile transcripts. To prioritize candidates, we computed Pearson correlation coefficients for the normalized GCLiPP profiles of 3′ UTRs of genes expressed in both Jurkat cells and primary mouse T cells (Figure 6A, black histogram) and examined transcript instability by RNAseq analysis of primary mouse T cells treated with actinomycin D (Figure 6A, red histogram). The proto-oncogene *PIM3* emerged as an outstanding candidate with both strong interspecies GCLiPP correlation and very high transcript instability. Alignment of the GCLiPP profiles of human and mouse *PIM3* revealed a dominant shared peak of GCLiPP read density (Figure 6B). This peak corresponded to a highly conserved region of the transcript that contains a G-quadruplex, followed by a putative AU-rich element (ARE) and a CELF binding motif (Figure 6C). Another conserved region with a G-quadruplex followed by a putative ARE appeared upstream of the biochemically shared GCLiPP peak. We numbered these conserved regions CR1 and CR2 according to their order in the 3′ UTR, and hypothesized that CR2 would exert greater cis-regulatory activity than CR1, given its RBP occupancy in both species and the relative lack of occupancy in CR1. To test this hypothesis, we performed CRISPR dissections of both the human and mouse *PIM3* 3′ UTRs (Figure 6—source data 1). These analyses produced largely concordant patterns of post-transcriptional cis-regulatory activity in the human and mouse 3′ with the greatest significant destabilizing effect corresponding to the shared region of GCLiPP read intensity covering the CR2 element (Figure 6D-K). Consistent with this portrait of the entire 3′ UTR, when we filtered specifically for mutations that completely deleted either CR1 or CR2, we observed significantly greater expression of transcripts derived from cells with CR2 deleted versus CR1 (Figure 6L, M). Thus, PIM3 is a very unstable transcript with highly concordant RBP occupancy in human and mouse. Functional dissection of the post-transcriptional regulatory landscape of this gene revealed that this biochemical concordance between mouse and human cells is mirrored at a functional level, with the most highly occupied region indicated by GCLiPP read density corresponding to the most destabilizing region of the 3′ UTR.

**Figure 6.**
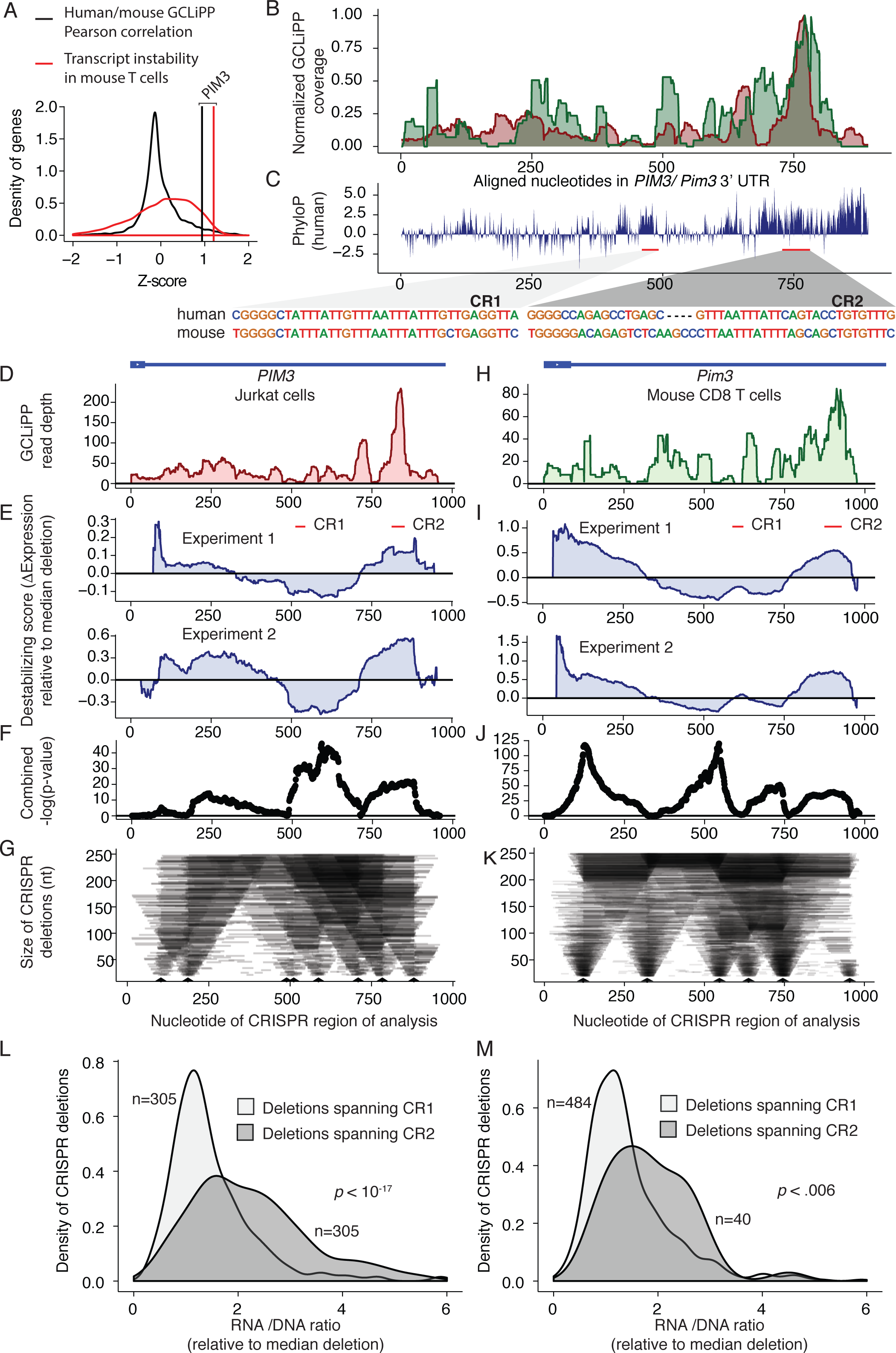
Biochemically and functionally shared post-transcriptional regulation of PIM3 in human and mouse cells. **(A)** Z-scores of Pearson correlation between mouse and human GCLiPP (black distribution) and transcript instability as measured by comparing transcript read abundance in untreated versus actinomycin treated mouse T cells (red distribution) for 7541 genes with matched data. Vertical lines indicate observations for PIM3. (**B**) Normalized human and mouse GCLiPP read density and (**C**) PhyloP across aligned nucleotides of PIM3 3′ UTR (as depicted in Figure 5). Insets show sequences of putative regulatory elements. (**D-L**) Dissection of human PIM3 3′UTR in Jurkat T cells (**D**) GCLiPP peaks aligned to schematic illustration of 3′UTR. (**E**) Change in expression along the 3′UTR relative to median expression of all possible deletions. Per-nucleotide effect score was calculated by comparing median normalized RNA/gDNA ratio for all shown deletions spanning a given nucleotide with all shown deletions. Experiment 1 and 2 are biological duplicates which were transfected with 80µM or 120µM of gRNAs respectively. Red bars indicate putative ARE-containing cis regulatory elements. (**F**) Unadjusted -log10 p-values from Welch’s two sample t-test comparing all deletions spanning a nucleotide with all other deletions across both experiments. (**G**) Size of deletions generated using CRISPR-Cas9. Arrow heads represent gRNA placement. (**H-K**) Dissection of mouse PIM3 3′UTR. Data are represented identically to human data, except that mouse primary CD8 T cells were used, and both mouse experiments 1 and 2 used a gRNA concentration of 80µM. (**L**) Effect of deletions spanning putative ARE containing cis-regulatory elements. The RNA/DNA ratio for mutants deleting ARE1 and ARE2 are shown in human Jurkat T cells. (**M**) Same as in (**L**) but using data from mouse primary T cells.

## Discussion

Interconnected networks of bound RBPs and RNAs form a complex layer of post-transcriptional regulation that affects all biological processes. Understanding these networks remains one of the key challenges in deciphering how the genome encodes diverse cell identities and behaviors. The outcomes of RNA/RBP interactions can be quite varied. RBP occupancy can affect RNA biogenesis, decay, translation, localization, splicing, chemical modification and editing. Developing a roadmap to understand the cis-regulatory elements in each gene will be critical to full elucidation of the post-transcriptional biology of any given transcript. Biochemical procedures like CLIP that utilize RBP-specific immunoprecipitation have facilitated decoding of these networks for individual RBPs. However, the number of validated CLIP antibodies remains much smaller than the number of RBPs which are known to associate with mature RNAs (Baltz et al., 2012; Sundararaman et al., 2016). In addition, the relative occupancy of cis-regulatory elements bound by different proteins cannot be directly compared using different pools of immunoprecipitated material. Here, we adapted previously described methodologies to arrive at a technique, GCLiPP, to provide such a global RBP occupancy roadmap, and applied this technique to interrogate post-transcriptional cis-regulatory activity in human and mouse T cells.

Systematic comparison with eCLIP data for 87 individual RBPs (Sundararaman et al., 2016) indicated that GCLiPP roughly represented a weighted average of all potential eCLIP experiments for cytosolic RBPs. Presumably, this includes proteins that are not appreciated as having RNA binding activity, and those for which no specific affinity reagents are currently available. GCLiPP peaks overlapped eCLIP peaks at a frequency much greater than would be expected by chance, and overall GCLiPP read density correlated with eCLIP read density in a manner that corresponded with the relative abundance of a given RBP in purified cellular mRNPs (Baltz et al., 2012). Nevertheless, the eCLIP peaks for some low abundance RBPs were significantly enriched in GCLiPP profiles. The strongest correlations were observed for abundant cytosolic RBPs, and the correspondence between eCLIP and GCLiPP was only apparent for cytosolic, but not non-cytosolic, RBPs. This result was expected. COMPARTMENTS annotations (Binder et al., 2014) indicate that most of the RBPs classified as non-cytosolic are mainly located in the nucleus, and the GCLiPP protocol includes pelleting nuclei after a gentle detergent based cellular lysis, followed by an enrichment for polyadenylated RNA. Both of these steps would be predicted to selectively eliminate nuclear RNPs associated with primary transcripts. Future iterations of GCLiPP could be modified to intentionally enrich for nuclear RBPs to examine the regulatory landscape of mRNA biogenesis.

The GCLiPP datasets described here provide a rich resource for the annotation and experimental dissection of cis-regulatory function in mRNAs. GCLiPP detected RBP occupancy at many known cis-regulatory regions, including canonical polyadenylation signals and elements that control mRNA localization, translation and stability, providing a biochemical correlate of functional activity. In addition, we demonstrated that GCLiPP can guide discovery of novel cis-regulatory elements. Dissection of the *CD69* 3′UTR revealed regions of destabilizing activity that correspond to RBP occupied sites detected by GCLiPP. Our findings corroborate a previous study that identified region PR2 as a potential destabilizing region that contained AREs (Santis et al., 1995). We expanded on this work and further identified a new destabilizing region (region PR1) that also exhibits GCLiPP evidence of RBP occupancy. Further assessment of these post-transcriptional cis-regulatory regions may provide novel insights into immune cell biology. CD69 is an important regulator of T cell differentiation (Martín et al., 2010), migration (Shiow et al., 2006) and metabolism (Cibrian et al., 2016). Because of these effects, CD69 is considered a potential target of therapeutic treatments for autoimmune and inflammatory disorders (González-Amaro et al., 2013). Generation of post-transcriptionally modified T cells expressing high levels of CD69 could provide insight into its role in T cell biology and inflammatory disorders.

We leveraged the matched datasets from similar cell types expressing many shared transcripts to perform a cross species comparison of the post-transcriptional regulatory landscape. As might be expected, the sequences of 3′ UTR regions that appeared as peaks of RBP occupancy in both species were in general more conserved than the full length 3′ UTRs in which they occurred. These biochemically shared peaks were enriched in well-known RBP-binding cis-regulatory sequences including PUM motifs, CELF motifs and canonical polyadenylation signals. Surprisingly, though, we also found clear biochemically shared peaks with relatively poor sequence conservation. These regions retain RBP occupancy despite an evident lack of strong selective pressure on their primary sequence, perhaps due to highly degenerate and/or structural determinants of RBP occupancy. RNAs with conserved structure and RBP binding but poorly conserved primary sequence have been reported before, and they are enriched in gene regulatory regions (Seemann et al., 2017; Weinreb et al., 2016). Finally, we noted that transcripts with biochemically shared peaks tended to encode proteins that were themselves involved in post-transcriptional gene regulation. This pattern is consistent with previous suggestions that auto-regulatory or multi-component feedback loops may be a conserved mode of post-transcriptional gene regulation (Kanitz and Gerber, 2010).

Our dissection of the human *PIM3* and mouse *Pim3* 3′UTRs demonstrates the utility of GCLiPP for decoding biochemically shared and functionally conserved post-transcriptional regulation. The PIM family of serine/threonine kinases exert profound regulatory effects on MYC activity, cap-dependent translation independent of MTOR, and BAD mediated antagonism of apoptosis (Narlik-Grassow et al., 2014). Post-transcriptional regulation of PIM kinases is important, as proviral integrations in the *Pim1* 3′ UTR are highly oncogenic (Nawijn et al., 2011). *Pim3* mRNA was abundant but highly labile in T cells, with a turnover rate in the top 2% of expressed mRNAs. PIM family members contain multiple ARE like repeats of AUUU(A), but the specific sequences responsible for rapid mRNA decay have not been described and cannot be predicted from the primary sequence alone. The *PIM3* 3′UTR contains two phylogenetically conserved regions with very similar predicted ARE sequences. Of these regions, we predicted that greater regulatory activity would be exerted by the region with GCLiPP evidence for RBP occupancy in both human and mouse cells. CRISPR dissection bore out this prediction in both species. The inactive conserved region may be structurally inaccessible to RBP occupancy, or it may be occupied and exert regulatory activity only in other cell types or signaling conditions. These possibilities further highlight the utility of unbiased biochemical determination of RBP occupancy for annotating the regulatory transcriptome. The datasets reported here will accelerate the annotation of cis-regulatory elements operant in T cell transcripts. In general, GCLiPP can be combined with other unbiased biochemical assays and genetic analyses to yield a roadmap for the dissection of post-transcriptional regulatory networks.

## Materials and Methods

### Cells

Jurkat cells were grown in RPMI supplemented with fetal bovine serum (Omega). Primary CD4^+^ and CD8^+^ mouse T cells were isolated from C57BL/6J mouse peripheral lymph nodes and spleen using positive and negative selection Dynabeads, respectively, according to the manufacturer’s instructions (Invitrogen). All mice were housed and bred in specific pathogen-free conditions in the Animal Barrier Facility at the University of California, San Francisco. Animal experiments were approved by the Institutional Animal Care and Use Committee of the University of California, San Francisco. Cells were stimulated with immobilized biotinylated anti-CD3 (clone 2C11, 0.25 µg/mL, BioXcell) and anti-CD28 (clone 37.51, 1 µg/mL, BioXcell) bound to Corning 10 cm cell culture dishes coated with Neutravidin (Thermo) at 10 µg/mL in PBS for 3 h at 37 °C. Cells were left on stimulation for 3 days before being transferred to non-coated dishes in T cell medium (Steiner et al., 2011) supplemented with recombinant human IL-2 (20 U/mL). Th2 cell cultures were also supplemented with murine IL-4 (100 U/mL) and anti-mouse IFN-g (10 µg/mL). CD8 T cell cultures were also supplemented with 10 ng/mL recombinant murine IL-12 (10 ng/mL). For re-stimulation, cells were treated with 20 nM phorbol 12-myristate 13-acetate (PMA) and 1 µM ionomycin (Sigma) for 4 hours before harvest.

### Measurement of mRNA Decay

Cells were stimulated with PMA and Ionomycin for 4 hours and then additionally treated with Actinomycin-D (Sigma-Aldrich) at 5 µg/mL for an additional 0, 1, 2 or 4 hours. After treatment, cells were lysed with Trizol LS (Life Technologies) and processed with Direct-zol ™ 96 well RNA (Zymogen). RNA was quantified with an ND-1000 spectrophotometer (NanoDrop) and reverse transcribed with SuperScript III First Strand Synthesis Kit (Invitrogen). Quantitative PCR was performed in two separate experiments using SYBR Advantage qPCR Premix (Clontech) on a Realplex 2S instrument (Eppendorf).

### GCLiPP and RNAseq

~100 × 10^6^ mouse T cells cultured from 3 mice or ~100 × 10^6^ Jurkat T cells were washed and resuspended in ice cold PBS and UV irradiated with a 254 nanometer UV crosslinker (Stratagene) in three doses of 4000 mJ, 2000 mJ and 2000 mJ, swirling on ice between doses. Cells were pelleted and frozen at -80 °C. Thawed pellets were rapidly resuspended in 400 µL PXL buffer without SDS (1X PBS with 0.5% deoxycholate, 0.5% NP-40, Protease inhibitor cocktail) supplemented with 2000 U RNasin (Promega) and 10 U DNase (Invitrogen). Pellets were incubated at 37 °C with shaking for 10 min, before pelleting of nuclei and cell debris (17000 g for 5 min). Supernatants were biotinylated by mixing at room temperature for 30 min with 500 µL of 10 mM EZ-Link NHS-SS-Biotin (Thermo) and 100 µL of 1 M sodium bicarbonate. Supernatants were mixed with 1 mg of washed oligo-dT beads (New England Biolabs) at room temperature for 30 min and washed 3 times with magnetic separation. Oligo-dT selected RNA was eluted from beads by heating in poly-A elution buffer (NEB) at 65 °C with vigorous shaking for 10 min. An aliquot of eluted RNA was treated with proteinase K and saved for RNAseq analysis using Illumina TruSeq Stranded Total RNA Library Prep Kit according to the manufacturer’s instructions. Cells treated with Actinomycin-D as described above were also collected for RNAseq to generate transcriptome wide measurements of transcript stability.

The remaining crosslinked, biotinylated mRNA-RBP complexes were captured on 250 µL of washed M-280 Streptavidin Dynabeads (Invitrogen) for 30 min at 4 °C with continuous rotation to mix. Beads were washed 3 times with PBS and resuspended in 40 µL of PBS containing 1000 U of RNase T1 (Thermo) for 1 min at room temperature. RNase activity was stopped by addition of concentrated (10% w/v) SDS to a final concentration of 1% SDS. Beads were washed successively in 1X PXL buffer, 5X PXL buffer and twice in PBS. 24 pmol of 3′ radiolabeled RNA linker was ligated to RBP bound RNA fragments by resuspending beads in 20 µL ligation buffer containing 10 U T4 RNA Ligase 1 (New England Biolabs) with 20% PEG 8000 at 37 degrees for 3 h. Beads were washed 3X with PBS and free 5′ RNA ends were phosphorylated with polynucleotide kinase (New England Biolabs). Beads were washed 3X with PBS and resuspended in ligation buffer containing 10 U T4 RNA Ligase 1, 50 pmol of 5′ RNA linker and 20% PEG 8000 and incubated at 15 °C overnight with intermittent mixing. Beads were again washed 3 times in PBS and linker ligated RBP binding fragments were eluted by treatment with proteinase K in 20 µL PBS with high speed shaking at 55 °C. Beads and supernatant were mixed 1:1 with bromophenol blue formamide RNA gel loading dye (Thermo) and loaded onto a 15% TBE-Urea denaturing polyacrylamide gel (BioRad). Ligated products with insert were visualized by autoradiography and compared to a control ligation (19 and 24 nt markers). Gel slices were crushed and soaked in gel diffusion buffer (0.5 M ammonium acetate; 10 mM magnesium acetate; 1 mM EDTA, pH 8.0; 0.1% SDS) at 37 °C for 30 min with high speed shaking, ethanol precipitated and resuspended in 20 µL of RNase free water. Ligated RNAs were reverse transcribed with Superscript III reverse transcriptase (Invitrogen) and amplified with Q5 polymerase (New England Biolabs). PCR was monitored using a real time PCR thermal cycler and amplification was discontinued when it ceased to amplify linearly. PCR products were run on a 10% TBE polyacrylamide gel, size selected for an amplicon with the predicted 20-50 bp insert size to exclude linker dimers, and purified from the gel (Qiagen). Cleaned up library DNA was quantified on an Agilent 2100 Bioanalyzer using the High Sensitivity DNA Kit before being sequenced. All GCLiPP and RNAseq sequencing runs were carried out on an Illumina HiSeq 2500 sequencer.

### GCLiPP and RNAseq bioinformatics analysis pipeline

FastQ files were de-multiplexed and trimmed of adapters. Each experiment was performed on three technical replicates per condition (resting and stimulated) per experiment. Cloning replicates and experiments were pooled in subsequent analyses. Jurkat and mouse T cell trimmed sequence reads were aligned to the hg38 human or mm10 mouse genome assembly using bowtie2, respectively. After alignment, PCR amplification artifacts were removed by de-duplication using the 2-nt random sequence at the 5′ end of the 3′ linker using a custom script that counted only a single read containing a unique linker sequence and start and end position of alignment per sequenced sample. Peaks of GCLiPP read density were called by convolving a normal distribution against a sliding window of the observed read distribution. A 70 nucleotide window was analyzed centered on every nucleotide within the 3′ UTR. For each window, the observed distribution of read density was compared to a normal distribution of the same magnitude as the nucleotide in the center of the window. The Pearson’s correlation coefficient was computed for each nucleotide and peaks were defined as local maxima of goodness of fit between observed GCLiPP read density and the normal distribution, requiring a read depth above 20% of the maximum read depth in the 3′ UTR global minimum of 10 reads. RNAseq reads were aligned using STAR Aligner (https://github.com/alexdobin/STAR) (Dobin et al., 2013) to align against the mm10 genome, and gene expression data were calculated as fragments per kilobase per million reads. All custom scripts are available as STAR Methods Key Resource.

### Comparison of GCLiPP to individual eCLIP datasets

eCLIP data (Sundararaman et al., 2016) were downloaded via the ENCODE data portal (http://www.encodeproject.org/). The first replicate set of bigwig files were downloaded for each RBP deposited online at the time of analysis (December 2017) as well as CLIPper called peaks for the same. To facilitate comparisons with GCLiPP we called GCLiPP peaks in the Jurkat data using CLIPper (Lovci et al., 2013) after re-aligning Jurkat GCLiPP reads to hg19. Correlation analysis was performed with a custom perl script that calculated the Spearman correlation for read depth at each nucleotide in the 3′ UTR of all genes that were expressed in each dataset (as determined by CLIP read depth). ~5000-15000 expressed genes were included in the correlation analysis for each RBP. For comparison to mRNP abundancy, log10 RBP mass spectrometry spectra counts were utilized from (Baltz et al., 2012). To stratify RBPs by subcellular localization, data were taken from the COMPARTMENTS database, with RBPs with a localization score of 5 in the cytosol counted as cytosolic and lower counted as non-cytosolic (Binder et al., 2014). All custom scripts are available as STAR Methods Key Resource.

### CRISPR editing

Guide RNA sequences were selected using the Benchling online CRISPR design tool (https://benchling.com/crispr) with guides selected to target genomic regions of GCLiPP read density. Synthetic crRNAs and tracrRNA (Dharmacon) were resuspended in water at 160 µM at 1:1 ratio and allowed to hybridize at 37 c for 30 m. For CRISPR dissection experiments, all crRNAs were mixed at an equimolar ratio before annealing to tracrRNA. This annealed gRNA complex (80 µM) was then mixed 1:1 by volume with 40 µM *S. pyogenes* Cas9-NLS (University of California Berkeley QB3 Macrolab) to a final concentration of 20 µM Cas9 ribonucleotide complex (RNP). This complexed gRNA:Cas9 RNP was mixed with a carrier solution of salmon sperm DNA (Invitrogen) and diluted to a final concentration between 5-20 µM. The diluted gRNA:Cas9 RNPs were nucleofected into primary mouse T cells (24 hours after stimulation) with the P3 Primary Cell 96-well Nucleofector ™ Kit and into Jurkat cells with the SE Cell Line 96-well Nucleofector ™ Kit using a 4-D Nucleofector following the manufacturer’s recommendations (Lonza). Cells were pipetted into pre-warmed media and then returned to CD3/CD28 stimulation for another two days and expanded an additional 3 days (mouse primary T cells) or cultured for 7-10 days (Jurkat).

### Quantification of CD69 protein and mRNA

Jurkat cells were gene edited using the CRISPR-Cas9 system as described above. After 3 days in culture, cells were washed and stained with anti-human CD69 PE (BioLegend, clone FN50). Samples were acquired on a FACSAria II (BD Biosciences) with CD69^hi^ cells single cell sorted into a 96 well plate and incubated at 37°C at 5% CO_2_ to allow generation of single cell clones. To verify editing, gDNA was extracted from Jurkat clones using QuickExtract ™ DNA Solution (Epicentre) according to the manufacturer’s instructions. Samples were then amplified by PCR using Q5^®^ High Fidelity DNA Polymerase (New England Biolabs). Quantitative PCR reactions were performed on an Eppendorf Realplex 2S thermocycler with the following program: (95°C 60 s; 35 cycles of 95°C 30 s, 58°C 30 s, 72°C 60 s).

### 3′ UTR dissection

3′ UTR dissection was performed as described (Zhao et al., 2017). Gene edited cells were harvested into Trizol reagent (Invitrogen) and total RNA was phase separated and purified from the aqueous phase using the Direct-zol RNA miniprep kit with on-column DNase treatment (Zymo). Genomic DNA was extracted from the remaining organic phase by vigorous mixing with back extraction buffer (4 M guanidine thiocyanate, 50 mM sodium citrate, 1 M Tris base). cDNA was prepared with oligo-dT using the SuperScript III reverse transcription kit (Invitrogen). cDNA and genomic DNA were used as a template for PCR using MyTaq 2X Red Mix (Bioline). To equilibrate the number of target molecules and number of PCR cycles between samples, we performed semi-quantitative PCR followed by agarose gel electrophoresis to determine a PCR cycle number where genomic DNA first showed visible bands. This cycle number was then used with a titration of cDNA concentrations and a concentration that amplified equivalently was selected for analysis by deep sequencing. To quantify relative RNA/DNA ratios, cDNA and genomic DNA amplicons were purified using a QIAquick PCR purification up kit (Qiagen) and quantified on an Agilent 2100 Bioanalyzer using the High Sensitivity DNA Kit (Agilent).

Amplicons were tagmented with the Nextera XT kit (Illumina) and sequenced on an Illumina 2500 HiSeq. Reads were aligned to a custom genome consisting of the targeted PCR amplicon using STAR aligner and mutations were scored using an awk script (https://github.com/alexdobin/STAR/blob/master/extras/scripts/sjFromSAMcollapseUandM.awk). RNA/DNA read ratios were calculated for all mutations over 20 nucleotides long and less than 250 nucleotides long, and relative expression was quantified as the median normalized RNA/DNA ratio for this subset of mutations. Mutations had to have at least 10 reads in both the RNA and gDNA amplicons and mutations with an RNA/DNA ratio of greater than 10 were excluded as outliers. Effect sizes for each nucleotide of the amplicon in each experiment were computed by comparing this median normalized RNA/DNA ratio for all mutations spanning a given nucleotide to all other mutations. Combined p-values were calculated using a Welch’s two sample t-test comparing all mutations spanning a given nucleotide with all other mutations.

### Shared peak calling, motif analysis and icSHAPE and Phylogenetic analyses

3′ UTR alignments of mouse and human were performed by downloading hg38 RefSeq 3′ UTRs from UCSC genome browser, (http://genome.ucsc.edu), identifying syntenic regions of the mouse genome in mm10 with the KentUtils liftOver program (https://github.com/ucscGenomeBrowser/kent) and aligning UTRs with Clustal Omega (http://www.ebi.ac.uk/Tools/msa/clustalo/) (Sievers et al., 2011). Biochemically shared peaks were called by the following algorithm: Measure normalized GCLiPP read density (i.e. the fraction of the maximal read depth within that 3′ UTR) at each position. Calculate correlation between mouse and human normalized signal, as well as observed data and a normal distribution centered at the point being examined in both the mouse and human data tracks. These three Spearman correlations were added together to calculate a numerical score, and shared peaks were defined as local maxima of these scores. To identify high stringency peaks, peaks were only accepted if they 1) had a correlation of >0.75 between mouse and human, 2) had a peak that had a read density of >0.5 of the maximum read density within that 3′ UTR in one data track (mouse or human) and >0.2 in the other and 3) had >10 reads at that location in both mouse and human datasets. Biological enrichment of genes with shared peaks was calculated using the Metascape (Tripathi et al., 2015) online interface (http://metascape.org) using the default settings, with the exception that a background set of genes was included in the analysis, specifically all genes that contain a called GCLiPP peak in both human and mouse datasets that do not contain a biochemically shared peak.

For motif calling, HOMER (Heinz et al., 2010) was used in RNA mode with the “noweight” option to turn off GC correction to search for motifs of width 5, 6 or 7 nucleotides, with otherwise default parameters. The positive sequence set was the mouse and human sequences of the biochemically shared GCLiPP peaks, the negative sequence set was all other GCLiPP called peaks from Jurkat and mouse T cells that were not shared across species. For icSHAPE we used a published bigwig file of locally normalized icSHAPE signal intensity generated in mouse ES cell (Spitale et al., 2015). Conservation of loci in the mouse and human genomes were obtained from the UCSC genome browser as a bigwig of PhyloP scores of conservation across 60 placental mammals (mouse) and 100 vertebrates (human) (http://hgdownload.cse.ucsc.edu/goldenpath/mm10/phyloP60way/, http://hgdownload.cse.ucsc.edu/goldenpath/hg38/phyloP100way/).

Oligonucleotide and primer sequences

GCLiPP 3′ RNA linker: 5′-NNGUGUCUUUACACAGCUACGGCGUCG-3′

GCLiPP 5′ RNA linker: 5′-CGACCAGCAUCGACUCAGAAG-3′

GCLiPP Reverse transcription primer: 5′-

CAAGCAGAAGACGGCATACGAGATNNNNNNCGCTAGTGACTGGAGTTCAGACGTGT

GCTCTTCCGATCCGACGCCGTAGCTGTGTAAA-3′ (NNNNNN is barcode for demultiplexing)

GCLiPP 3′ PCR primer: 5′-CAAGCAGAAGACGGCATACGAGAT-3′

GCLiPP 5′ PCR primer: 5′-

AATGATACGGCGACCACCGAGATCTACACTGGTACTCCGACCAGCATCGACTCAGAAG-3′

Read1seq sequencing primer for GCLiPP: 5′-

ACACTGGTACTCCGACCAGCATCGACTCAGAAG-3′Index sequencer primer for GCLiPP: 5′-GATCGGAAGAGCACACGTCTGAACTCCAGTCAC-3′

CD69 gRNA1: CTCAAGGAAATCTGTGTCAG

CD69 gRNA2: TCATTCTTGGGCATGGTTAT

CD69 gRNA3: CCTGTGATGCTTCTAGCTCA

CD69 gRNA4: AATAATGAAATAACTAGGCG

CD69 gRNA5: TAATTGAATCCCTTAAACTC

CD69 gRNA6: TGATGTGGCAAATCTCTATT

PIM3 (human) gRNA1: TGTGCAGGCATCGCAGATGG

PIM3 (human) gRNA2: GACTTTGTACAGTCTGCTTG

PIM3 (human) gRNA3: GTGGCTAACTTAAGGGGAGT

PIM3 (human) gRNA4: AAACAATAAATAGCCCCGGT

PIM3 (human) gRNA5: TTGAGAAAACCAAGTCCCGC

PIM3 (human) gRNA6: CAGGAGGAGACGGCCCACGC

PIM3 (human) gRNA7: TTTATGGTGTGACCCCCTGG

PIM3 (human) gRNA8: CCAAGCCCCAGGGGACAGTG

Pim3 (mouse) gRNA1: GTTCAATTCTGGGAGAGCGC

Pim3 (mouse) gRNA2 CTGGTTCAAGTATCCACCCA

Pim3 (mouse) gRNA3: CCATAAATAAGAGACCGTGG

Pim3 (mouse) gRNA4: GCTTCCTCCCGCAAACACGG

Pim3 (mouse) gRNA5: CTGGTGTGACTAAGCATCAG

Pim3 (mouse) gRNA6: TGGAGAAGGTGGTTGCTTGG

Primers

CD69 F: TGGAATGTGAGAAGAATTTATACTGG

CD69 R: GTAATAGAATTGATTTAGGAAAG

PIM3 F (human): TCCAGCAGCGAGAGCTTGTGAGGAG

PIM3 R(human): TGATCTCCAGACATCTCACTTTTGAACTG

PIM3 R2(human):

TGAGATAGGTGCCTCACTGATTAAGCATTGGTGATCTCCAGACATCTCACTTTTGAACTG

Pim3 F (mouse): GCGTTCCAGAGAACTGTGACCTTCG

Pim3 R (mouse): TATGATCTTCAGACATTTCACACTTTTG

## Author Contributions

Author Contributions: A.L. and W.S.Z. performed the experiments and performed bioinformatic analyses. R.K. established the bioinformatic pipeline for small RNA sequencing analysis. W.Z. and D.E. helped design CRISPR dissection experiments. N.Z. consulted on data analysis and interpretation. K.M.A., A.L. and W.S.Z. designed experiments, interpreted the data, and wrote the manuscript. All authors discussed the results and approved the manuscript.

## Data availability

Datasets in this paper are available on Gene Expression Omnibus accessions GSE94 and GSE115886

## Acknowledgements

We thank David Siegel for advice on analyzing pooled crRNP 3′ UTR dissection experiments. A.L. was supported by Cancer Research Institute Irvington Fellowship and the UCSF Immunology T32 training grant T32AI007334. This work was supported by the US National Institutes of Health (HL107202, HL109102, AI128047, HL124285, GM110251), the Sandler Asthma Basic Research Center, and a Scholar Award (K.M.A.) from The Leukemia & Lymphoma Society. The authors declare no conflicts of interest.

